# An elongated COI fragment to discriminate botryllid species and as an improved ascidian DNA barcode

**DOI:** 10.1101/2021.01.20.427448

**Authors:** Marika Salonna, Fabio Gasparini, Dorothée Huchon, Federica Montesanto, Michal Haddas-Sasson, Merrick Ekins, Marissa McNamara, Francesco Mastrototaro, Carmela Gissi

## Abstract

Botryllids are colonial ascidians widely studied for their potential invasiveness and as model organisms, however the morphological description and discrimination of these species is very problematic, leading to frequent specimen misidentifications.

To facilitate species discrimination and detection of cryptic/new species, we developed new barcoding primers for the amplification of a COI fragment of about 860 bp (860-COI), which is an extension of the common Folmer’s barcode region. Our 860-COI was successfully amplified in 177 worldwide-sampled botryllid colonies. Combined with morphological analyses, 860-COI allowed not only discriminating known species, but also identifying undescribed and cryptic species, resurrecting old species currently in synonymy, and proposing the assignment of clade D of the model organism *Botryllus schlosseri* to *Botryllus renierii*. Importantly, within clade A of *B. schlosseri*, 860-COI recognized at least two candidate species against only one recognized by the Folmer’s fragment, underlining the need of further genetic investigations on this clade. This result also suggests that the 860-COI could have a greater ability to diagnose cryptic/new species than the Folmer’s fragment at very short evolutionary distances, such as those observed within clade A. Finally, our new primers simplify the amplification of 860-COI even in non-botryllid ascidians, suggesting their wider usefulness in ascidians.

## INTRODUCTION

The subfamily Botryllinae (family Styelidae) consists of small colonial ascidians of the genera *Botryllus* and *Botrylloides*. It includes species widely studied because of their invasive potential, ecological importance, or as model organisms in developmental biology and for investigating processes such as regeneration, stem cells migration, apoptosis and allorecognition ^1–6^. Despite their importance, the taxonomy of Botryllinae species is highly debated, since their small zooids show few hardly visible discriminant characters. In addition, colony growth occurs by a peculiar process of asexual reproduction by palleal budding, in which the anatomical structures of each zooid are continuously developed and reabsorbed, so most morphological features exhibit high intraspecific variability ^7,8^. Therefore, poor and often confusing descriptions of the botryllid species have accumulated over time, with synonymies or invalid descriptions of new species frequently proposed (see the WoRMS, Ascidiacea World Database, at http://www.marinespecies.org/ascidiacea/) and cryptic species identified in widely studied species ^9^. Even the diagnostic characters distinguishing the genera *Botrylloides* and *Botryllus* have been amended several times (see discussion in ^10^), with the validity of the genus *Botrylloides* rejected in 1987 ^11^. Currently, the most accepted Botryllinae taxonomy considers both genera as valid according to the definition of Milne Edwards ^12^, with a single diagnostic character consisting of the presence (in *Botryllus*) or absence (in *Botrylloides*) of an atrial siphon, and resulting in structural differences of the common cloaca ^10^.

The difficulties of Botryllinae morphological species description and identification demonstrate the need for new discriminant characters, to complement the classical morphological ones.

In 2001, Saito proposed new criteria for Botryllinae species classification and phylogeny such as the examination of the life history, the type of sexual reproduction, the vascular system formation, and the allorecognition process ^13,14^. Although very intriguing, a taxonomy based on these criteria is often impossible, since it requires the observation of living colonies, sometimes for extended periods, which obviously cannot be applied to museum specimens. Indeed, these new classificatory criteria have been so far successfully applied only to the identification and description of some Japanese species ^14–17^ and to the reconstruction of an integrated phylogeny of few Japanese botryllids (in combination with 18S rDNA and classical morphological data) ^13^.

DNA sequences are another category of data useful for species description and discrimination as well as for phylogenetic inferences. So far, a 500-600 bp fragment of the mitochondrial cytochrome oxidase subunit I (COI) gene has been used in several phylogenetic studies on *Botryllus schlosseri* and Styelidae, as well as for botryllid species discrimination ^9,18–20^, *i.e.,* as DNA barcode ^21^. This fragment has been amplified mainly with Folmer’s primers ^22^ (so it is hereafter named “Folmer’s fragment”), although the effectiveness of these primers in ascidians is not very high, as also testified by the numerous published alternative primers ^23–28^. The 18S rDNA, the mitochondrial cytochrome *b* (*cob*) gene, and the entire mitochondrial genome (mtDNA) have been also used to try to elucidate the relationships within the subfamily Botryllinae ^13,29,30^ and within the family Styelidae ^19^, or to detect botryllid cryptic species ^31^. Most of these studies have left the overall Botryllinae phylogeny poorly resolved, because of the low support of several nodes in the reconstructed trees ^19,29^, but they have been decisive for discriminating species and detecting cryptic speciation events. Indeed, both nuclear and mitochondrial markers have demonstrated the weaknesses of the *Botryllus schlosseri* morphological taxonomy, revealing that *B. schlosseri* is a “species complex” consisting of five genetically highly divergent clades (named from A to E), each corresponding to a cryptic species and with a peculiar geographic distribution ^9,18–20,32,33^. *B. schlosseri* is a model organism widely used for investigating fundamental biological processes (such as regeneration, stem cells migration, apoptosis and allorecognition) ^3,5,6^ but it is also a cosmopolitan fouling species and a widespread marine invaders ^9,34–36^. Therefore, it is crucial to fully resolve its taxonomic status.

Recently, we have successfully applied to Botryllinae an integrative taxonomy approach combining morphological characters and sequence data ^37,38^. In particular, the analysis of the entire mtDNA and of the zooid/colony morphology has allowed us to recognize clade E as a new species, *Botryllus gaiae* ^38^, distinct from the *B. schlosseri* clade A, for which an accurately described neotype was previously designated ^39^. In another study, the morphological characters have been analysed together with an elongated COI fragment of about 860 bp (hereafter named 860-COI), leading to a better description and an effective discrimination of three *Botrylloides* species having highly similar appearance ^37^.

In order to set a better molecular tool for botryllid identification and for the detection of cryptic or new species, we here evaluate the use of new barcoding primers and of the related 860-COI fragment for the discrimination of 177 worldwide-sampled botryllid colonies, including archival specimens and specimens with uncertain taxonomic assignment or with only a gross ‘in field’ morphological characterization. Specifically, we have firstly compared *in silico* the effectiveness of the new primers for 860-COI to that of the Folmer’s primers and performed PCR experiments to verify the successful rate of our primers. Then, through tree reconstructions and species delimitation analyses, we have compared the ability of the 860-COI and the Folmer’s fragment to discriminate known botryllid species, and to identify cryptic/candidate new species for which further clues are available in the literature. Whenever possible, specimens of the candidate species have been also morphologically analysed. Finally, we have tested the 860-COI primers in ascidians other than botryllids, using both PCR and *in silico* comparisons: the results lead to the proposal that 860-COI could be used as DNA barcode for most ascidian taxa.

## METHODS

### Botryllid specimens

A total of 134 new botryllid colonies were analysed in this study in conjunction with 43 additional botryllid samples whose 860-COI fragment was published in our previous studies ^37,38,40^. The specimen collection is described in detail in Supplementary Table S1, which reports sampling localities (from Europe to Australia, Japan, California and Brazil), collection dates, initial and final species assignment, and other useful information. This collection includes specimens from several source: from archival specimens of museum institutions, to colonies kindly provided by ascidian specialists, up to *ad hoc* or fortuitous samplings. For some specimens, a formalin-preserved fraction of the same colony was also available. As reported in Supplementary Table S1, among the archival specimens there are: the holotype and paratype of *Botrylloides conchyliatus* (Queensland Museum), the paratype of *Botrylloides crystallinus* ^40^, two syntypes of *Botryllus gaiae* ^38^, nine Australian *Botrylloides* without a clear assignment at species level (Queensland Museum, QM, Australia), and two colonies of *Botryllus eilatensis* (the Steinhardt Museum of Natural History, Israel). As for the last species, it will be reported hereafter as *Botrylloides eilatensis*. Indeed, *Botryllus eilatensis* is characterized by the absence of an atrial siphon and a consequent peculiar structure of the common cloaca and the zooid systems ^41^, all characters distinctive of the genus *Botrylloides*, as described by ^10,12^. Therefore, this species should be moved from the genus *Botryllus* to the genus *Botrylloides* and we propose the revised binomial of *Botrylloides eilatensis.* Six specimens assigned only at genus level come from the monitoring of the Western Australia coasts, while colonies with a trustworthy assignment at species level were provided by the ascidian taxonomists Y. Saito (17 Japanese samples) and R.M. Rocha (three Brazilian samples). Finally, nine *Botryllus* colonies come from a demersal fauna study with trawl net in the Ionian Sea (Italian coasts: station 72 near Tricase, and station 74 near Porto Badisco, see Supplementary File S1). Although the formalin-preserved material of these specimens was in poor conditions, some specific morphological features (*i.e.,* the shape of the colony, the thickness of the matrix, and the stomach and testis peculiarities) rule out its belonging to *B. schlosseri sensu* Brunetti ^39^ and indicate a high similarity to *Botryllus renierii* (Lamarck, 1815) as described in Brunetti ^42^ (see a detailed morphological description of these specimens in Supplementary File S1). Therefore, these *Botryllus* specimens will be reported hereafter as “putative *B. renierii”*, although their bad condition suggests the application of a precautionary approach, and a consequent final identification as “*Botryllus* indet”. Indeed, according to the Open Nomenclature qualifiers ^43^, “indet” means that the specimen is indeterminable beyond a certain taxonomic level due to the deterioration or lack of diagnostic characters.

As for the *B. schlosseri* species complex *sensu* Bock ^9^, 68 specimens come from *ad hoc* periodic samplings in the Venice Lagoon (North Adriatic Sea, Italy), aimed at obtaining colonies for the *B. schlosseri* breeding laboratory of Padua University. These colonies were collected at the same site near Chioggia, at a depth of 1-2 m, and found on *Zostera* leaves, ropes or floats. Likewise, an *ad hoc* sampling of 15 *B. schlosseri* colonies was performed in Vilanova (Spain), since this is one of the few localities where the rare B and C clades were previously found ^9,18^. Other *B. schlosseri* specimens from California and France were provided by researchers specifically working on this model organism. Finally, 11 colonies of the *B. schlosseri* species complex were fortuitously found in South Italy during samplings with methods/gears unusual for ascidians and devoted to:

- clam sampling with a dredge in the Adriatic Sea, near Barletta, at 2.5-4 m depth (5 colonies);
- benthic fauna study by scuba diving in the Mar Piccolo and Mar Grande of the Gulf of Taranto, Ionian Sea, at 2-3 m depth (6 colonies).

### Primers for 860-COI

The 860-COI fragment was amplified with the primer pair dinF/Nux1R or according to a nested-PCR strategy using the dinF/Nux1R primers in combination with the nested primers cat1F and ux1R (Table 1 and Figure 1-A). The latter strategy was carried out only when the PCR with dinF/Nux1R resulted in low product yield or no visible amplicon on agarose gel. In this case, a 1:100 dilution of the first amplification was directly used as template of the second PCR with the primer pair cat1F/ux1R.

**Table 1.**
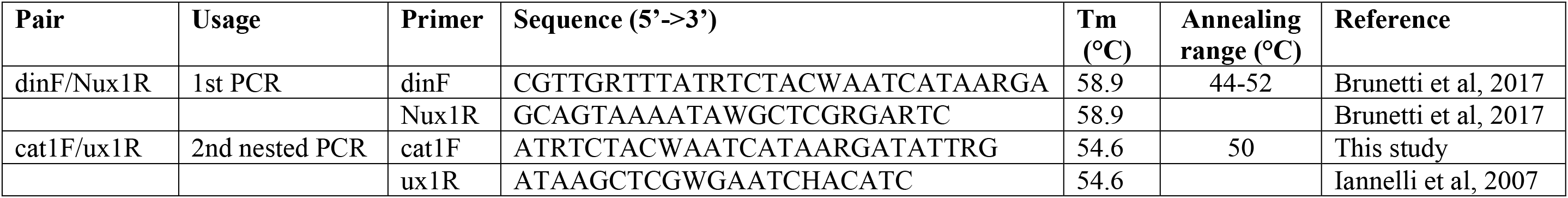
Primer pairs used in this study

**Figure 1.**
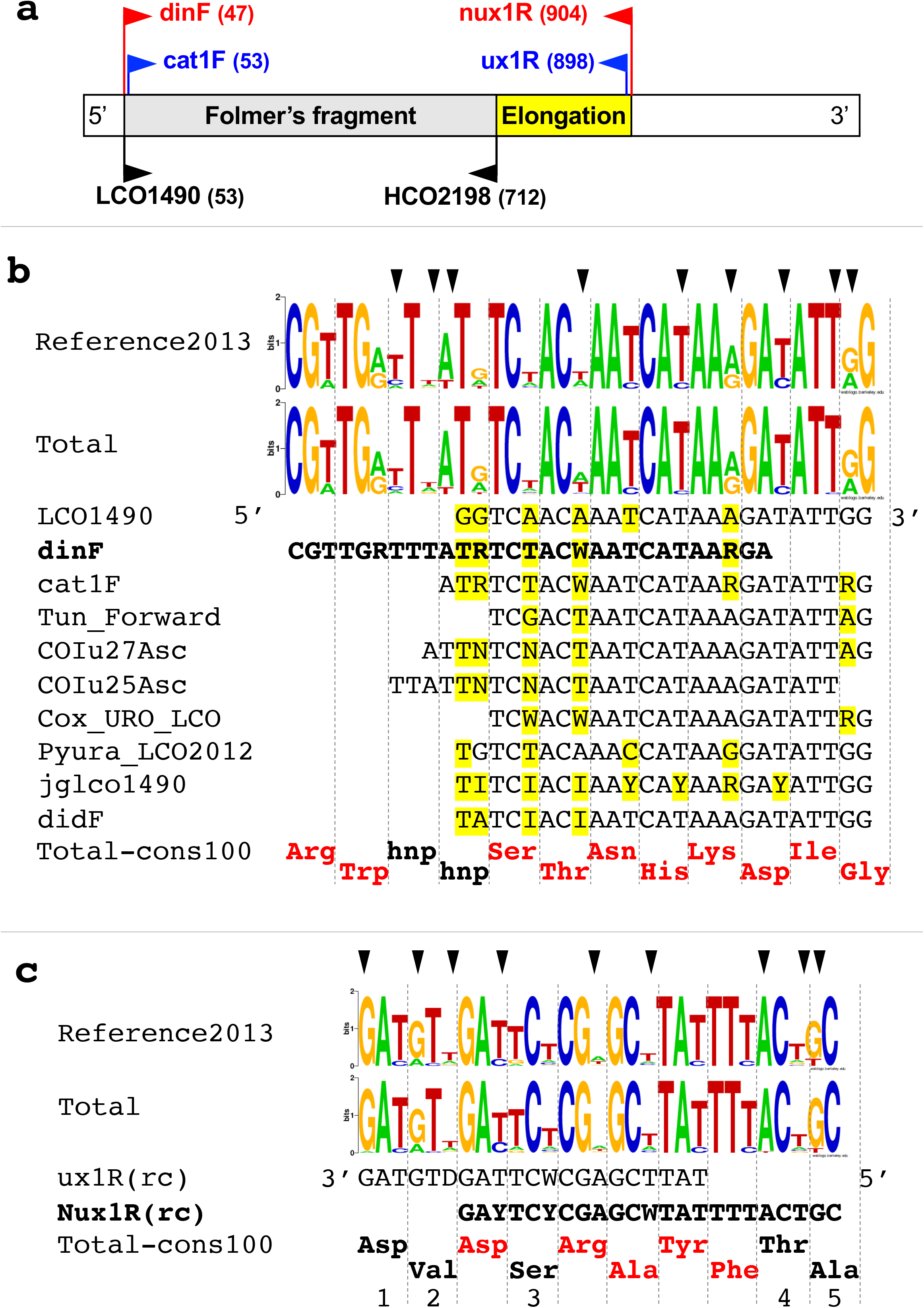
Mapping and description of the primers for the 860-COI fragment a) Primer mapping on the COI of *B. schlosseri*. Red: primers for the first PCR; blue: nested primers for the second (reamplification) PCR; in brackets: position of the primer 3’ end on the complete COI sequence of *B. schlosseri* (1548 bp, Acc. number: FM177702). b) Forward primers aligned to the ascidian COI. Reference2013: WebLogo plot of the dataset of 18 complete COI sequences of ascidians used for primer design; Total: WebLogo plot of the dataset of 33 complete COI sequences of ascidians currently available; Total-cons100: 100% amino acid consensus sequence from the Total dataset; yellow: positions with differences to the LCO1490 primer; red amino acids: positions with a 100% conservation; hnp: the hydrophobic non-polar amino acids Met, Leu, Phe, or Val; arrow head: positions differing between the two logos. Sequence datasets are reported in Supplementary Table S2. Primer references: Cox_URO_LCO and Pyura_LCO2012 are described in Rubinstein ^27^; COIu27Asc and COIu25Asc in Monniot ^24^; Tun_Forward in Stefaniak ^28^; didF in da Silva ^92^; jglco1490 in Geller ^71^. c) Our reverse primers aligned to the ascidian COI. Rc: primer in reverse-complement orientation. All other symbols are as in b). Differences from the amino acid consensus sequence are as below: 1= Asn in *Lissoclinum patella* (KJ596323), with A in the 1st codon position; 2= Thr in *Diplosoma listerianum* (FN313539); Ile in 6 species of Aplousobranchia and Styelidae; 3= Ala in two *Phallusia* species; Thr in *Lissoclinum patella* (KJ596323), *Diplosoma listerianum* (FN313539) and *Didemnum vexillum* (KM259616); 4= Ser in *Lissoclinum patella* (KJ596323); 5= Ser in three *Ciona* species

These primers were designed by our group in 2013 based on the analysis of the dataset “Reference2013”, consisting of the 18 complete COI sequences of ascidians available at that time, both as published and private data (see Supplementary Table S2). The additional 15 complete COI of ascidians made available from 2013 to 2019 (dataset “New2019” in Supplementary Table S2) were analysed *in silico* to evaluate primer effectiveness on other ascidian species. Both our primers and the Folmer’s primers (LCO1490 and HCO2198) were mapped on each COI sequence with the Primer3 software available in Geneious ver. 5.5.7.2 ^44^ using the options: maximum mismatches = 6; no mismatches within 2 bp of the 3’ end. In addition, the sequence conservation in the regions of primer mapping was represented with logo plots and consensus sequences. Sequence logo plots were made with WebLogo v. 2.8.2 (https://weblogo.berkeley.edu) ^45^ for both the Reference2013 dataset and the “Total” dataset, consisting of the Reference2013 plus the New2019 datasets. The strict consensus (threshold value of 100%) for the amino acid sequences of the “Total” dataset was calculated with Seaview ^46^.

### Primer design criteria

Figure 1 describes the amplification strategy of the 860-COI fragment, and compares the sequence and the position, along the COI gene, of our new primers to that of the LCO1490 and HCO2198 Folmer’s primers, as well as of other published primers mapping on the same region.

The forward primer dinF is an optimisation of the LCO1490 primer for ascidians, obtained considering some important but usually underestimated factors. Indeed, its sequence is based not only on the consensus sequence reconstructed from the Reference2013 dataset but takes also into account the overlooked degeneracy of the ascidian mt genetic code. In particular, the mismatched positions between LCO1490 and the consensus of the Reference2013 dataset have been modified in dinF in order to introduce in those positions either the most frequent residue or a degenerate base (Figure 1-B). Moreover, the 3’ end of the new dinF primer has been shifted 6 bp upstream compared to LCO1490, in order to ensure a better pairing to the template at the last and penultimate primer positions. Indeed, as shown in Figure 1-B, the Guanine (G) of the penultimate position of LCO1490 corresponds to a purine (R) in the Reference2013 dataset, since it falls on the 1st codon position of a conserved Gly. This situation stems from the peculiarities of the ascidian mt genetic code, where Gly is encoded not only by the GGN (as in the universal code) but also by the AGR codons ^47,48^, thus both A and G can be present at the 1st position of the Gly codons. The possible mispairing of LCO1490 at the 3’ penultimate position could be responsible of the efficiency reduction of this primer in ascidians (Figure 1-B; see Discussion and references therein), so in our new primer we avoid this position by shifting the 3’ end of dinF upstream, that is at level of the 1st+2nd codon position of the conserved and negatively charged amino acid Asp (GAY - with Y= pyrimidines- in Reference2013 of Figure 1-B). In this way a perfect primer-template pairing should take place at the last and penultimate positions of dinF (*i.e.,* GA) even in case of conservative substitutions at the Asp site, since the two negatively charged amino acids, Asp and Glu, are globally encoded by GAN codons. The nested forward primer cat1F was designed in order to have exactly the same boundaries of LCO1490 but includes the degeneracies and base changes introduced in dinF (Figure 1-B).

To increase the number of COI sites usable in phylogenetic analyses, the reverse primer Nux1R has been positioned about 200 bp downstream of the annealing region of HCO2198 (Figure 1-A). Specifically, Nux1R has been designed by improving the ascidian-specificity of our old primer ux1R, that worked quite well in amplifying the mtDNA of several ascidian species ^31,49^. Even in this case, the Reference 2013 dataset has guided the identification of the primer positions to be degenerated or set to the most abundant base, and the degeneracy on the ascidian mt genetic code was considered for the definition of the 3’ terminus of Nux1R (Figure 1-C).

### DNA extraction

Most botryllid colonies were preserved in 99% ethanol, and a few were stored in RNAlater at −20°C. Total DNA was extracted from a colony fragment including the tunic or from few isolated zooids, mainly according to a modified CTAB method ^50^. The NucleoSpin DNA RapidLyse Kit (Macherey-Nagel) was used for the DNA extraction from the Queensland Museum archival specimen QM_G335164. For non-botryllid ascidians (Table 2 and Supplementary Table S3), total DNA was extracted from isolated zooids, gonads or siphonal muscles depending on the species, using the protocol detailed in ^51^ or the previously cited modified CTAB method ^50^. The usage of different DNA extraction methods/kits was related only to the different laboratory in which it was carried out, not to specimen features.

**Table 2.**
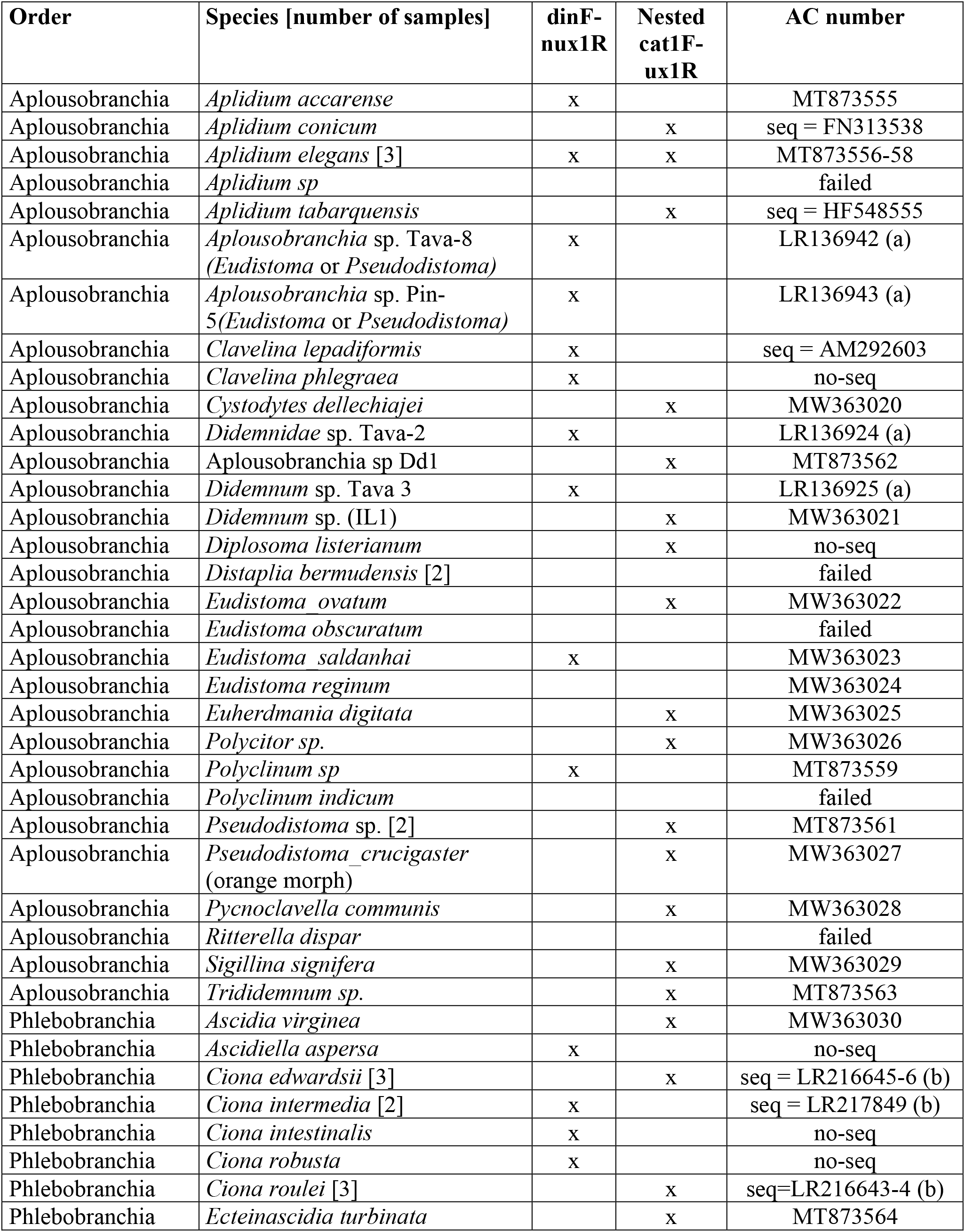
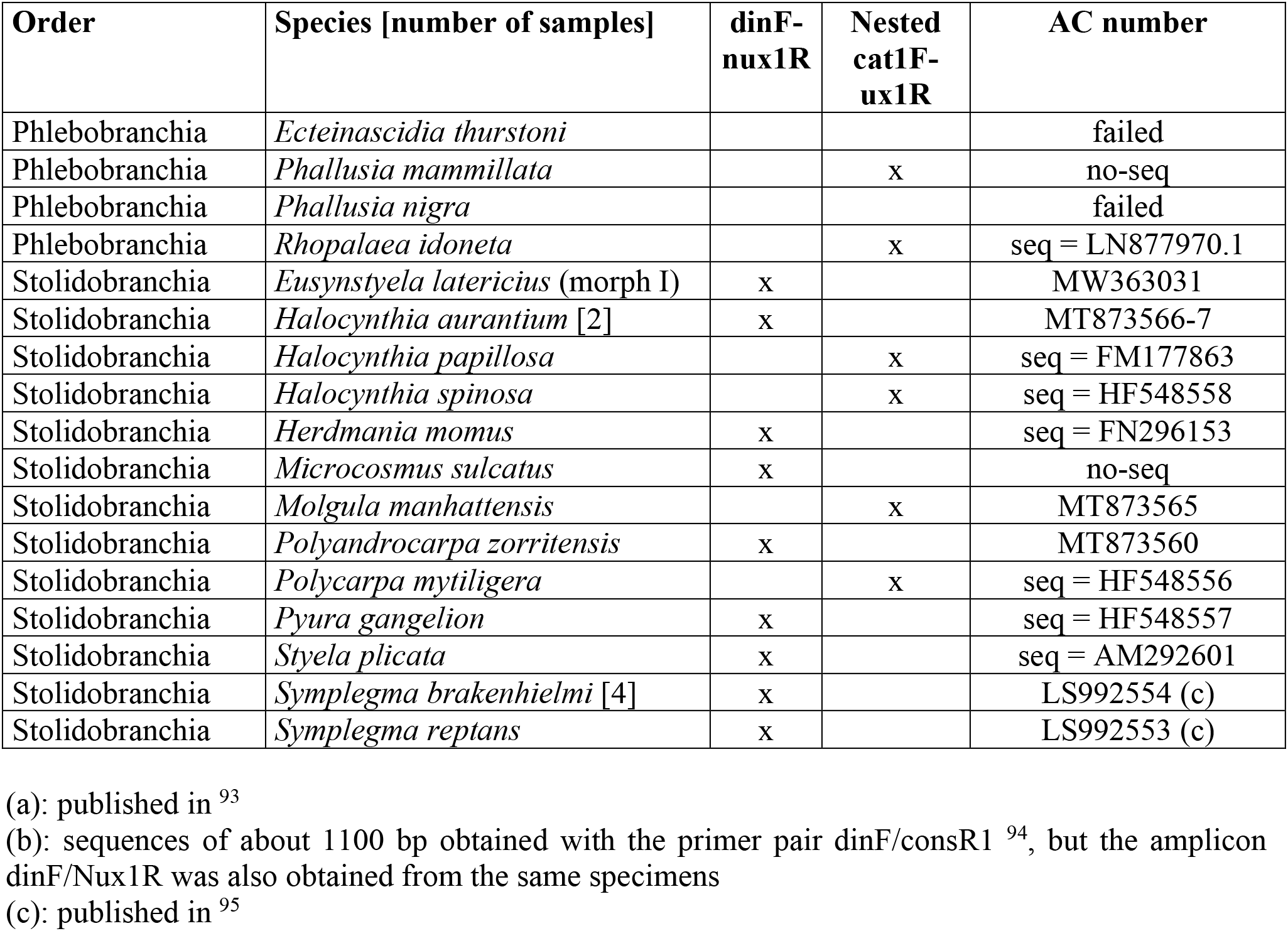
Amplification of the 860-COI in non-botryllid ascidian species. “no-seq”: amplicon checked by gel electrophoresis but not sequenced; “seq=”: sequenced amplicon, producing a sequence equal to that of an AC number already in the ENA nucleotide database

### Amplification of 860-COI

Depending on the laboratory, the 860-COI amplification reactions were accomplished with high-fidelity (Ex Taq, PrimeStar HS or PrimeStar GXL DNA polymerases of the TaKaRa Bio Inc.) or with standard (DreamTaq - Thermo Fisher Scientific, or EmeraldAmp Max HS PCR Master Mix - TaKaRa Bio Inc) DNA polymerases. The annealing temperature was in the range of 44-52°C for the primer pair dinF/Nux1R, and 50°C for the primers cat1F/ux1R (Table 1). Details on the amplification conditions are reported in Supplementary File S2.

All sequences were deposited at the ENA or GenBank nucleotide databases (see AC numbers in Supplementary Table S1 and in Table 2)

### Sequence analyses

The 860-COI sequences obtained from the 177 colonies clustered in 49 distinct haplotypes (= different sequences; see Supplementary Table S1) and were analysed together with the public COI sequences of botryllids and *Symplegma* listed in Supplementary Table S4 (data from the NCBI nt-nr database at Nov 2019). In order to exclude bias due to substantial differences in sequence length, we only included in the alignment sequences longer than 800 bp, with the exception of four short (524-671 bp) sequences representatives of the *B. schlosseri* clade B and C, for which no longest sequences were available in the public databases. Thus, the aligned dataset consists of a total of 55 Botryllinae and two *Symplegma* sequences.

Our dataset also contains public sequences of *Botrylloides leachii* (Supplementary Table S4). According to Viard and coll. ^52^, all currently published COI sequences of *B. leachii* have to be re-assigned to *Botrylloides diegensis*, since they are identical to the COI sequences of new specimens claimed to be *B. diegensis.*. However, one should note that in Viard’s study the assignment of the new specimens to *B. diegensis* is based only on the colour pattern of the colonies, without description of the anatomical traits discriminating *B. leachii* from *B. diegensis* (*i.e.,* the number of rows of stigmata, the presence/absence of muscle in the branchial sac, and the number of stomach folds; see ^53^). In the absence of this morphological information, we chose to keep the original assignment to *B. leachii* of all previously published sequences as well as of the identical sequences obtained in this study, pending combined morphological and molecular analyses on additional specimens.

Sequences of *Symplegma* (subfamily Polyzoinae) were used as outgroups in the phylogenetic reconstructions, since previous morphological and molecular data placed this genus sister to botryllids^34,54^.

Sequences were translated and aligned at amino acid level with MAFFT v 7. ^55^. The alignment was then manually optimized, and reverse translated to the corresponding nucleotide (nt) alignment. The obtained alignment consisted of a total of 1548 nt sites, since it included also several complete COI sequences (see Supplementary Table S4). However, it was not used in the successive analyses due to the length heterogeneity of the single sequences and then to the presence of many gapped sites. Therefore, starting from this long alignment, two shortest alignments were built and used in phylogenetic reconstructions and species delimitation analyses, that is:

- the “Elongated-856nt” alignment, containing all 856 sites obtainable by amplification with our primers, and thus corresponding to the elongated 860-COI fragment;
- the “Folmer-524nt” alignment, containing only the 524 sites obtained by amplification with the Folmer’s primers or with other primers matching at the same positions. This alignment consists of the same sites/region analysed in previous phylogenetic studies on botryllids ^9,18,20,32,56,57^.

Phylogenetic trees were reconstructed with the Maximum Likelihood (ML) method and by Bayesian Inference (BI). For ML, we used the online PhyML-SMS v3.0 software, which includes the automatic model selection algorithm Smart Model Selection (SMS) ^58,59^ (http://www.atgc-montpellier.fr/phyml-sms/). The best-fit substitution model selected under the Akaike Information Criterion (AIC) was the GTR+I+G for both the Elongated-856nt and the Folmer-524nt alignments. The proportion of invariant sites (I) and the gamma shape parameter (alpha) for the 4 rate categories were estimated by the PHYML v3.0 software itself. Bootstrap values, indicating node reliability, were based on 100 replicates. BI were performed with MrBayes v. 3.2.7a ^60^ under the GTR+I+G model, *i.e.*, the model selected by PhyML-SMS. Two parallel analyses, each composed of one cold and three incrementally heated chains, were run for 1,000,000 generations. Trees were sampled every 100 generations and the results of the initial 250,000 generations were discarded (burn-in fraction of 25%), after verifying that stationarity of the lnL was reached. As convergence diagnostic, we verified, at the end of the run that the average standard deviation of split frequencies was below 0.01 and that the PSRFs (Potential Scale Reduction Factor) were always close to 1.0 according to the indications reported in the MrBayes manual. Therefore, a total of 7,500 trees were used to calculate the Bayesian posterior probabilities (BPP) at the different nodes.

Species delimitation analyses were carried out on both the “Elongated-856nt” and the “Folmer-524nt” alignments using two different approaches: the Automatic Barcode Gap Discovery method (ABGD) ^61^ which is a sequence similarity clustering method, and the Poisson Tree Processes (PTP) ^62^ which is a tree-based coalescence method. The hypothetical species identified by these methods will be hereafter referred as Operational Taxonomic Units (OTUs). ABGD clusters sequences into partitions, consisting of hypothetical species, based on the statistical inference of the “barcode gap”, *i.e.,* the gap in the distribution of intra-species and inter-species pairwise distances. As demonstrated by tests on real data, the ABGD initial partitions are generally stable and identify OTUs corresponding to the species described by taxonomists, while the recursive partitions better handle the sequence heterogeneity identifying a largest number of OTUs ^61^. PTP infers putative species boundaries on both ultrametric and non-ultrametric input trees assuming the existence of two independent classes of Poisson processes, one describing speciation and the other coalescent events. ABGD analyses ^61^ were performed on the web-based interface (http://wwwabi.snv.jussieu.fr/public/abgd/) (last accessed date: March 2020). Prior intraspecific divergence was settled from the value corresponding to a single nucleotide difference (Pmin=0.001) to the default value of 0.1 (Pmax). All other values were as default (Steps = 10, Nb bins = 20). Since the default value for the minimum relative gap width (X= 1.5) did not produce a result for either alignments, we used the highest values that gave ABGD results (*i.e.*, X=1.0 for the “Elongated-856nt” and X=1.1 for the “Folmer-524nt” alignment). All three metric options provided by ABGD for the pairwise distance calculations were used, that is: Jukes-Cantor (JC69) ^63^, Kimura 2 parameter (K80) ^64^ and the simple uncorrected p-distance (p-dist). This strategy allowed excluding possible bias of the selected evolutionary model on the OTU delimitation. Therefore, we carried out three ABGD analyses per alignment. For the PTP method ^62^, both the ML and the Bayesian trees reconstructed from each alignment were used as input, although PTP was demonstrated to be quite robust to the tree reconstruction method ^65^. PTP analyses were performed using the Bayesian implementation (bPTP) available on the web-based interface (http://species.h-its.org/ptp/) (last accessed date: March 2020). Analyses were performed after removing the outgroup species and using the following parameter values: 500,000 MCMC generations, thinning every 100 generations and a burn-in fraction of 0.30. The convergence of the MCMC chains was confirmed by visual inspection of the likelihood plot, according to the PTP manual (https://species.h-its.org/help/), and then the maximum likelihood solutions were recorded.

## RESULTS

### The 860-COI sequences of botryllids

Our new primers and nested PCR strategy (Table 1 and Figure 1) successfully amplified the elongated COI fragment in all 177 analysed Botryllinae specimens. The nested PCR was necessary in only 13 species/cryptic species, for a total of 33 specimens (Supplementary Table S1). Some specimens of the same species were successfully amplified just with the dinF/Nux1R pair or by nested PCR, thus indicating that DNA quality within species can vary enough to affect primer functioning. As shown in Table 3, our primers successfully worked in 15 *Botrylloides* species (total of 56 specimens) and six species/cryptic species of *Botryllus* (120 specimens), including four clades of the *B. schlosseri* species complex. Interestingly, the five COI haplotypes obtained from the nine “QM” specimens identified just as *Botrylloides* sp. turned out to be different species (see the “Species delimitation analyses” section) and were provisionally named from sp3 to sp7 (Table 3). Moreover, the 860-COI sequences of *Botryllus* indet., *i.e.,* the putative *B. renierii,* were highly similar and significantly clustered with the published COI sequences of clade D in all phylogenetic reconstructions (data not shown), thus supporting the hypothesis that clade D could be *B. renierii* (see also the “Finding rare clades” section).

**Table 3.** Botryllinae species characterized by the 860-COI fragment in this and in previous studies

| Species | Specimen N° |  | Haplotype N° |
| --- | --- | --- | --- |
|  | From this study | Already published |  |
| clade A sensu Bock et al. 2011 = *Botryllus schlosseri* sensu Brunetti 2017 | 88 | 4 (a) | 12 |
| clade C sensu Bock et al. 2011 | 2 |  | 1 |
| clade D sensu Bock et al. 2011 = *Botryllus* indet., putative *Botryllus renierii* | 9 |  | 2 |
| clade E sensu Bock et al. 2011 = *Botryllus gaiae* |  | 14 (b) | 7 |
| *Botryllus primigenus* | 2 |  | 2 |
| *Botryllus scalaris* | 1 |  | 1 |
| *Botrylloides conchyliatus* |  | 5 (c) | 2 |
| *Botrylloides crystallinus* | 1 |  | 1 |
| *Botrylloides eilatensis* (d) | 2 |  | 1 |
| *Botrylloides fuscus* | 6 | 1 | 1 |
| *Botrylloides giganteus* |  | 10 (c) | 3 |
| *Botrylloides leachii* (e) | 3 |  | 3 |
| *Botrylloides niger* | 5 |  | 2 |
| *Botrylloides perspicuus* |  | 8 (c) | 1 |
| *Botrylloides simodensis* | 5 | 1 (c) | 3 |
| *Botrylloides violaceus* | 1 |  | 2 |
| QM *Botrylloides* sp3 = *Botrylloides leptum* | 5 |  | 1 |
| QM *Botrylloides* sp4 = *Botrylloides* cf. *pannosum*: new species? | 1 |  | 1 |
| QM *Botrylloides* sp5 = new species | 1 |  | 1 |
| QM *Botrylloides* sp6 = *Botrylloides jacksonianum* | 1 |  | 1 |
| QM *Botrylloides* sp7 = *Botrylloides* cf. *anceps* | 1 |  | 1 |
| Total | 134 | 43 | 49 |

(a) in <sup>39</sup>. AC numbers: LT908007-10
(b) in <sup>38</sup>. AC numbers: LR745518, LR743459-65
(c) in <sup>37</sup>. AC numbers: HF922626, LS992550-2, LS992542-3, and LS992545-8.
(d) Binomial name revised from *Botryllus eilatensis* to *Botrylloides eilatensis*, since this species is characterized by the absence of an atrial siphon and a consequent peculiar structure of the common cloaca and the zooid systems <sup>41</sup>, all characters distinctive of the genus *Botrylloides* <sup>10,12</sup>.
(e) based on Viard et al. <sup>52</sup>, all published COI sequences of *B. leachii* should actually belong to *Botryllus diegensis*. This should hold also for our new sequences. Waiting for confirmation by more detailed morphological analyses on additional specimens, here we use the original taxonomic assignment.

### Primer effectiveness outside botryllids

Since our primers were designed on COI sequences belonging to all three ascidian orders (see dataset Reference2013 in Supplementary Table S2), we checked their effectiveness in non-botryllid ascidians, both *in silico* and experimentally. Thus, we evaluated *in silico* their pairing ability to the 33 complete COI of ascidians currently available, including those published after 2013 (see Supplementary Table S2). The comparison of the logo plots generated from the two datasets (i.e., Reference2013 and Total) highlights the presence of small differences in the relative base frequency only for few sites, mostly at the third codon position (see arrows heads in Figure 1-B and Figure 1-C). As detailed in Supplementary Table S2, most of the 33 complete COI sequences have less than 3 mismatches to our primers but 4-5 mismatches to the Folmer’s primers. In addition, one third of all analysed sequences show a mismatch at the penultimate 3’ end position of the Folmer’s LCO1490 forward primer (see “MP” in Supplementary Table S2). Therefore, in most species our primers, especially the forwards ones, should work better than the Folmer’s primers. However, ux1R may not work at all in some Aplousobranchia, since its last 3’ terminal position shows a mismatch to the complete COI of *Lissoclinum patella* and of a *Didemnum sp.* species (Huchon, confidential data), where an Asp to Asn amino acid substitution has been observed (see position “1” in Figure 1-C). As for the experimental checks, test PCRs were carried out taking advantage of 55 non-botryllid species whose total DNA was available in our laboratories: the 860-COI fragment was successfully amplified in all but seven species (Table 2). The few failed amplifications concern mainly Aplousobranchia species (Table 2), being these results in agreement with the *in silico* observation on our reverse primer ux1R (see position “1” in Figure 1-C). Overall, both *in silico* and experimental results suggest that our primers could work not only in botryllids, but in a wide ascidian taxonomic range.

### Sequence comparison

Excluding the two outgroup sequences, the percentage of variable sites is 44.16% in the “Elongated-856nt” alignment against 44.08% in the “Folmer-524nt” alignment and 43.83% in the “Elongation” region outside the Folmer’s fragment (*i.e.,* in the yellow region of Figure 1a). Thus, the level of variability is very similar in the two considered COI barcoding regions, with an almost uniform distribution of the variable sites in Folmer’s fragment and in the downstream “Elongation” region. Moreover, taking into account only the 12 very similar sequences of clade A, the percentage of variable sites becomes higher in the “Elongation” region than in the “Folmer-524nt” alignment (7.66% *versus* 5.53%).

Although the botryllid phylogeny was not the aim of our research, we reconstructed phylogenetic trees as a general method of sequence comparison and as prerequisite for the PTP species delimitation analyses. Thus, the topology of these reconstructed trees is here described. Figure 2 summarizes the results of the Bayesian and ML reconstructions performed on the alignment “Elongated-856nt”, *i.e*., the 860-COI fragment. Only some nodes are highly supported in both the ML and the Bayesian tree, and can therefore be considered reliably resolved (nodes with BPP ≥ 0.90 together with ML bootstrap ≥ 85% in Figure 2). These nodes identify all clades corresponding to known species (*i.e.*, clades containing specimens morphologically assigned to the same species), including the recently described *B. gaiae* (= the former *B. schlosseri* clade E) ^38^, and to cryptic species of the *B. schlosseri* species complex. Well resolved nodes also strongly support:

1. the sister relationship between *Botrylloides perspicuus* and *Botrylloides simodensis*;
2. the sister relationship between two unassigned *Botrylloides sp.* samples of the Queensland Museum, *i.e.*, QM_G335162 (*Botrylloides* sp 5) and QM_G335165 (*Botrylloides* sp 6);
3. a monophyletic group consisting of the former five clades (A-E) of the *B. schlosseri* species complex *sensu* Bock ^9^. This group should be now considered as a cluster of the genus *Botryllus,* since it consists of genetically identified cryptic species plus 2-3 well described species, *i.e., B. schlosseri sensu* Brunetti 2017 (=clade A) ^39^, *B. gaiae* (=the ex clade E) ^38^ and the putative *B. renierii* (=clade D);
4. the existence of three main sub-clades, named A1 to A3, within the *B. schlosseri* clade A (red dots in Figure 2, with the relative support values in Table 4). Noteworthy, sub-clade A2 includes the haplotype VE and the sequences of the topotype NeoB and NeoC, that have been associated to the morphological description of the recently designated *B. schlosseri* neotype ^39^. In addition, sub-clade A1 includes the sc6ab specimen, *i.e.* the specimen target of the *B. schlosseri* genome sequencing project ^66^.

Apart from few additional strongly supported nodes within clade A (Table 4), the remaining parts of the 860-COI tree remain unresolved (see nodes without support in Figure 2) or are reliably supported only by Bayesian inference (few basal nodes with BPP ≥ 0.90 in Figure 2). These results demonstrate that the 860-COI fragment efficiently discriminates between Botryllinae species and is also able to resolve relationships between very closely related specimens, *i.e.,* within the *B. schlosseri* clade A. However, in spite of the COI sequence elongation, 860-COI is unable to clarify the deepest phylogenetic relationships within Botryllinae.

**Figure 2.**
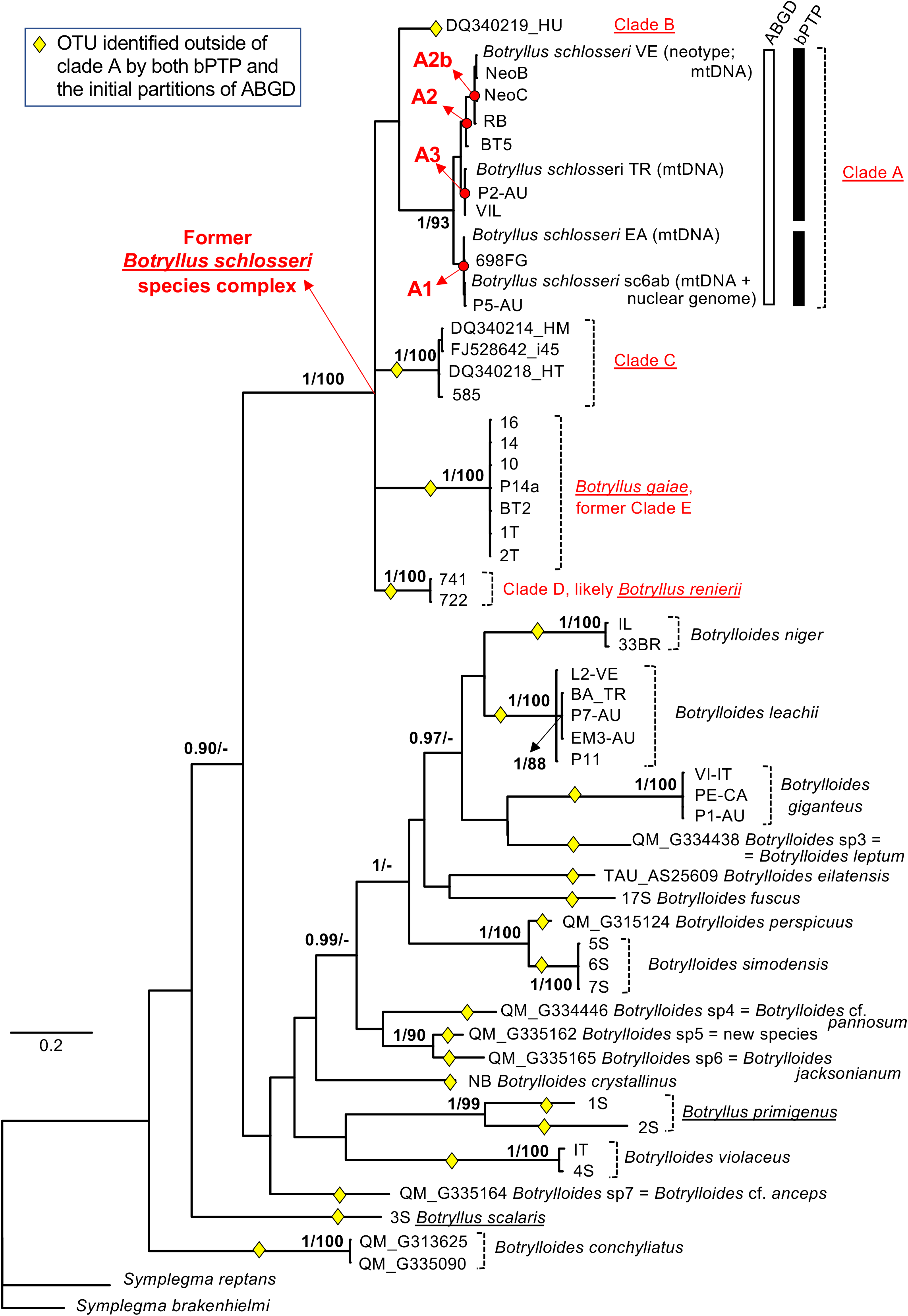
Bayesian majority rule consensus tree reconstructed from the “Elongated-856nt” alignment. Except for clade A, support values are reported close to the nodes as BPP/ML bootstrap percentage, only if BPP ≥ 0.90 or ML bootstrap ≥ 70%. Only for clade A, node supports are shown in Table 4, and the result of species delimitation analyses are reported as vertical bar (black for the ABGD initial partitions; white for bPTP). Yellow diamonds: OTUs identified outside of clade A by the initial partitions of all ABGD analyses (see Methods) and by bPTP; red dots: main sub-clades (A1, A2 and A3) within clade A, described in Table 4; underlined names: genus *Botryllus*. The specimens of *B. schlosseri* clade A used as neotype ^39^ and for which the entire mitochondrial or nuclear genomes were sequenced ^31,66^ are also indicated.

**Table 4.** Bayesian Posterior Probability (BPP) and ML bootstrap percentage for the nodes within clade A shown in Figure 2 and Figure 3. Red: BPP ≥ 0.90 or ML bootstrap ≥ 80%.

| Node | Alignment<br>Elongated-856nt |  | Alignment<br>Folmer-524nt |  |
| --- | --- | --- | --- | --- |
|  | BPP | ML boot. % | BPP | ML boot. % |
| A | 1.0 | 93 | 0.81 | 77 |
| A1 = (EA, sc6ab, P5-AU, 698FG) | 0.96 | 89 | 0.85 | 64 |
| A2 = (VE, NeoB, NeoC, RB, 5BT) =<br>(A2b, 5BT) | 0.94 | 95 | 0.68 | 66 |
| A2b = (VE, NeoB, NeoC, RB) | 0.99 | 100 | 0.98 | 97 |
| A3 = (TR, P2-AU, VIL) | 0.96 | 95 | 0.55 | 78 |
| (sc6ab, P5-AU) | 0.99 | 68 | 0.98 | 66 |
| (VE, NeoB) | 0.96 | 77 | 0.92 | 69 |
| (A2, A3) | 0.70 | 70 | <0.5 | <50 |

In order to verify if the elongated 860-COI fragment is more efficient to diagnose species compared to the shortest Folmer’s fragment, Bayesian and ML phylogenetic reconstructions have been carried out also on the shortest alignment “Folmer-524nt”. The resulting tree, shown in Figure 3, is very similar to that obtained from the Elongated-856nt alignment (Figure 2). The main difference is that the Folmer-524nt tree fails to support the sub-clades A1, A2 and A3, as well as other nodes within clade A, that are all well resolved by the Elongated-856nt alignment (Table 4). Similarly, the few basal nodes significantly supported in the Bayesian analysis of the Elongated-856nt alignment are unresolved in the Folmer-524nt Bayesian tree (compare Figure 2 to Figure 3). Overall, these results indicate that the 860-COI fragment behaves in general as the Folmer’s fragment and has a greater discrimination ability only within clade A of *B. schlosseri*, *i.e.*, at very short evolutionary distances. Of note, clade A is the only clade where the species delimitation analyses recognize a different number of OTUs/candidate species depending on the analyses alignment (“Elongated-856nt” or “Folmer-524nt”) and on the method (see the following paragraph).

**Figure 3.**
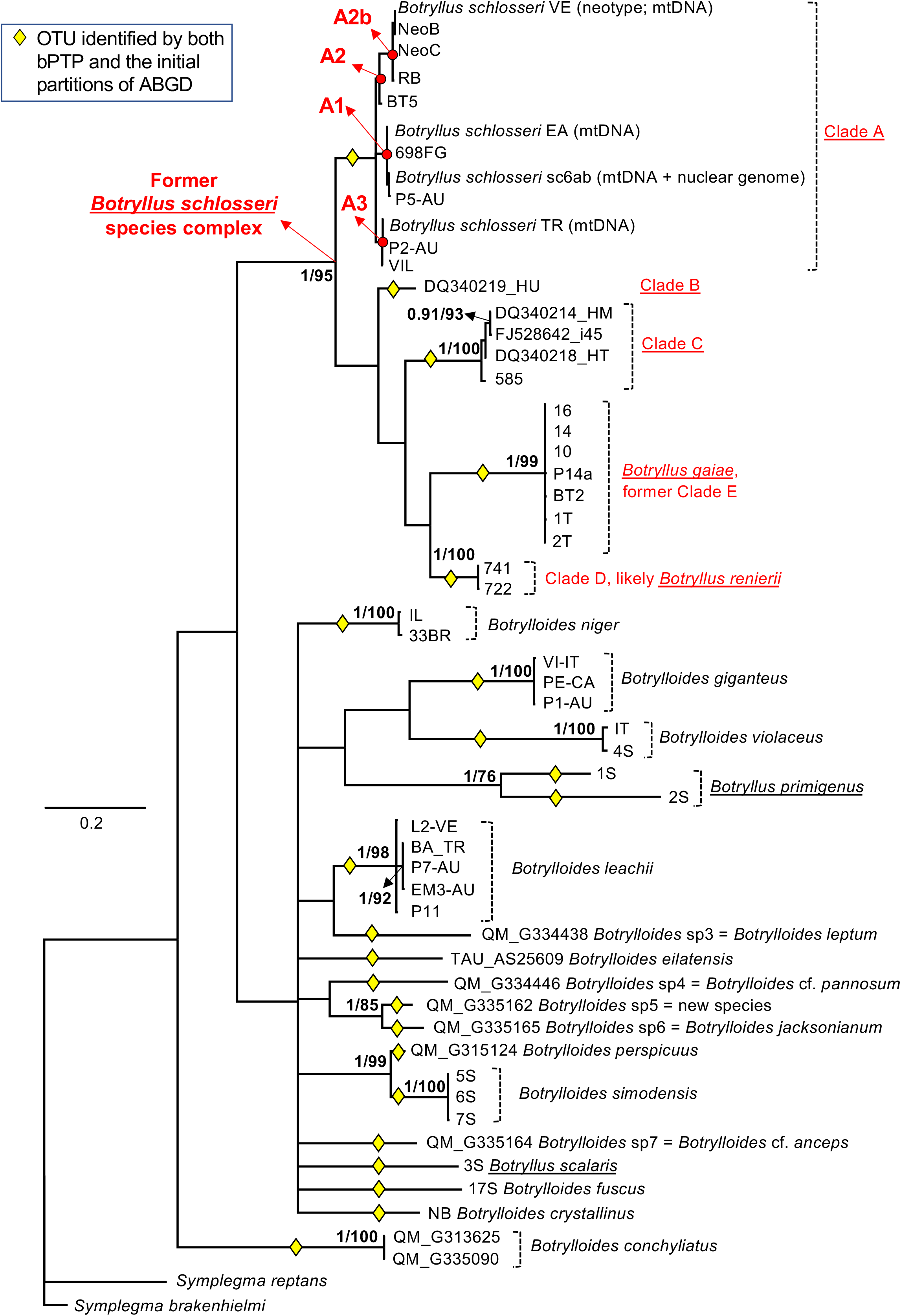
Bayesian majority rule consensus tree reconstructed from the “Folmer-524nt” alignment. Except for clade A, support values are reported close to the nodes as BPP/ML bootstrap percentage, only if BPP ≥ 0.90 or ML bootstrap ≥ 70%. Only for clade A, node supports are shown in Table 4. Yellow diamonds: OTUs identified by the initial partitions of all ABGD analyses (see Methods) and by bPTP; red dots: main sub-clades (A1, A2 and A3) within clade A, described in Table 4; underlined names: genus *Botryllus*. The specimens of *B. schlosseri* clade A as neotype ^39^ and for which the entire mitochondrial or nuclear genomes were sequenced ^31,66^ are also indicated.

### Species delimitation analyses

Outside clade A, the bPTP and the initial partitions of all ABGD analyses recognize exactly the same 22 OTUs in both the “Elongated-856nt” and the “Folmer-524nt” datasets (see yellow diamonds in Figure 2 and Figure 3). Each of these 22 OTUs corresponds to: a morphologically described species; one of the former clades of the *B. schlosseri* species complex (*i.e.* clade B, clade C, *B. gaiae* and *Botryllus* indet., corresponding to the putative *B. renierii*); one of the five different haplotypes coming from the QM specimens identified in the field just as *Botrylloides* sp (see Table 3).

Thanks to the availability of the corresponding formalin-preserved material, detailed morphological analyses have been carried out on the QM specimens in order to confirm the molecular result and to understand whether they belong to already described or to yet undescribed (*i.e.,* new) species. As reported in Table 3 and Supplementary Table S1, these morphological examinations have shown that:

- *Botrylloides* sp 3, *i.e.,* the OTU with the largest number of specimens, can be morphologically identified as *Botrylloides leptum* Herdman 1899;
- *Botrylloides* sp 4 (QM_G334446) is likely a new species similar to the original description of *Botrylloides pannosum* (Herdman, 1899), therefore for now it is reported as *Botrylloides* cf. *pannosum*;
- *Botrylloides* sp 5 (QM_G335162) is a stalked *Botrylloides* that cannot be assigned to any of the former described species, thus it is a new species;
- *Botrylloides* sp 6 (QM_G335165) is morphologically identified as *Botrylloides jacksonianum* (Herdman, 1899);
- *Botrylloides* sp 7 (QM_G335164) is another stalked *Botrylloides* and is morphologically very close to *Botrylloides anceps* (Herdman, 1891), therefore for now it is identified as *Botrylloides* cf. *anceps.*

Surprisingly, the two specimens of *Botryllus primigenus* are recognized by both methods not as a single but as two distinct OTUs, *i.e.,* as two different candidate species.

Within clade A, a different number of OTUs has been recognized depending on the dataset and the species delimitation method. Indeed, the bPTP on the “Elongated-856nt” dataset recognizes two OTUs within clade A, one corresponding to the sub-clade A1 and the other to the cluster A2+A3 (black vertical bar in Figure 2). On the contrary, the ABGD initial partitions on the “Elongated-856nt” dataset (white vertical bar in Figure 2) and the “Folmer-524nt” dataset (both bPTP and ABGD initial partitions; see yellow diamonds in Figure 3) recognize clade A as a single OTU. Differences in the type/number of OTUs identified within clade A are observed also in some ABGD recursive partitions (Table 5). Indeed, the recursive partitions perfectly match the initial partitions only at the highest values of prior intraspecific divergence, while at the lowest intraspecific values they recognize up to six new OTUs, depending on the dataset/distance metric (Table 5). In particular, these new OTUs always include sub-clade A1, while the remaining sequences of clade A are assembled in 1-3 additional OTUs depending on the different positioning of the 5BT specimen (thus, the additional OTUs can be: the single OTU (A2+A3); A2b plus the cluster (A3+5BT); or the three OTUs A2b, 5BT and A3; see Table 5).

**Table 5.** OTUs identified only in some recursive partitions of the ABGD analyses. OTU names are defined as in Table 4.

| Alignment | Distance | Prior intra-specific divergence |  | OTUs of the Recursive Partitions absent in the Initial Partitions |
| --- | --- | --- | --- | --- |
|  |  | From | To | OTU |
| Elongated-856nt | JC | 0.1 | 0.28 | A1; A2b; 5BT; A3 |
| " | K80 | 0.1 | 0.28 | A1; A2b; 5BT; A3 |
| " | p-dist | 0.1 | 0.46 | A1; (A2+A3) |
| Folmer-524nt | JC | 0.1 | 0.17 | A1; A2b; (A3+5BT);<br>*B. leachii* split in 2 OTUs;<br>*B. gaiae* split in 4 OTUs |
| " | " | 0.28 | 0.77 | A1; A2b; (A3+5BT) |
| " | K80 | 0.1 | 0.17 | A1; A2b; (A3+5BT);<br>*B. leachii* split in 2 OTUs;<br>*B. gaiae* split in 4 OTUs |
| " | " | 0.28 | 0.77 | A1; A2b; (A3+5BT) |
| " | p-dist | 0.1 | 0.28 | A1; A2b; 5BT; A3 |

Regarding the ABGD recursive partitions, it is noteworthy that the “Folmer-524nt” dataset recognizes a higher number of new OTUs than the “Elongated-856nt” dataset (Table 5), including also several OTUs within *B. leachii* and *B. gaiae* not supported by the current botryllid taxonomy or by our phylogenetic analyses of Figure 2 and Figure 3.

### Rare clades of B. schlosseri in the Mediterranean Sea

Our collection of the *B. schlosseri* clades *sensu* Bock ^9^ includes 113 specimens coming mainly from Mediterranean localities (Italian, French and Spanish coasts; see Supplementary Table S1). Although 78% of these specimens belong to the globally widespread clade A, our collection also provides interesting information on the spatio-temporal distribution of two rare clades/cryptic species in the Mediterranean Sea. Indeed, our non-systematic (and sometimes fortuitous) sampling reveals the following:

- the presence of the rare clade C in the Venice Lagoon in 2013 (2 over a total of 16 *B. schlosseri* colonies; see red triangle in Figure 4 and Supplementary Table S1);
- the presence of the rare clade D, a putative *B. renierii*, in two very close localities of the Ionian Sea, where they were detected during a demersal fauna study at a depth of 50-91 m (all 9 colonies sampled; see red star in Figure 4);
- the failure to find, thus the possible disappearance, of the rare clades B and C in Vilanova (Spain), *i.e.,* the only Mediterranean locality were clade C was found and the only worldwide locality of clade B ^9,18^. It should be noted that our Vilanova sampling was carried out in 2015 (15 colonies, all belonging to clade A; Supplementary Table S1), while the previous successful recoveries date back to 2005 ^18^.

**Figure 4.**
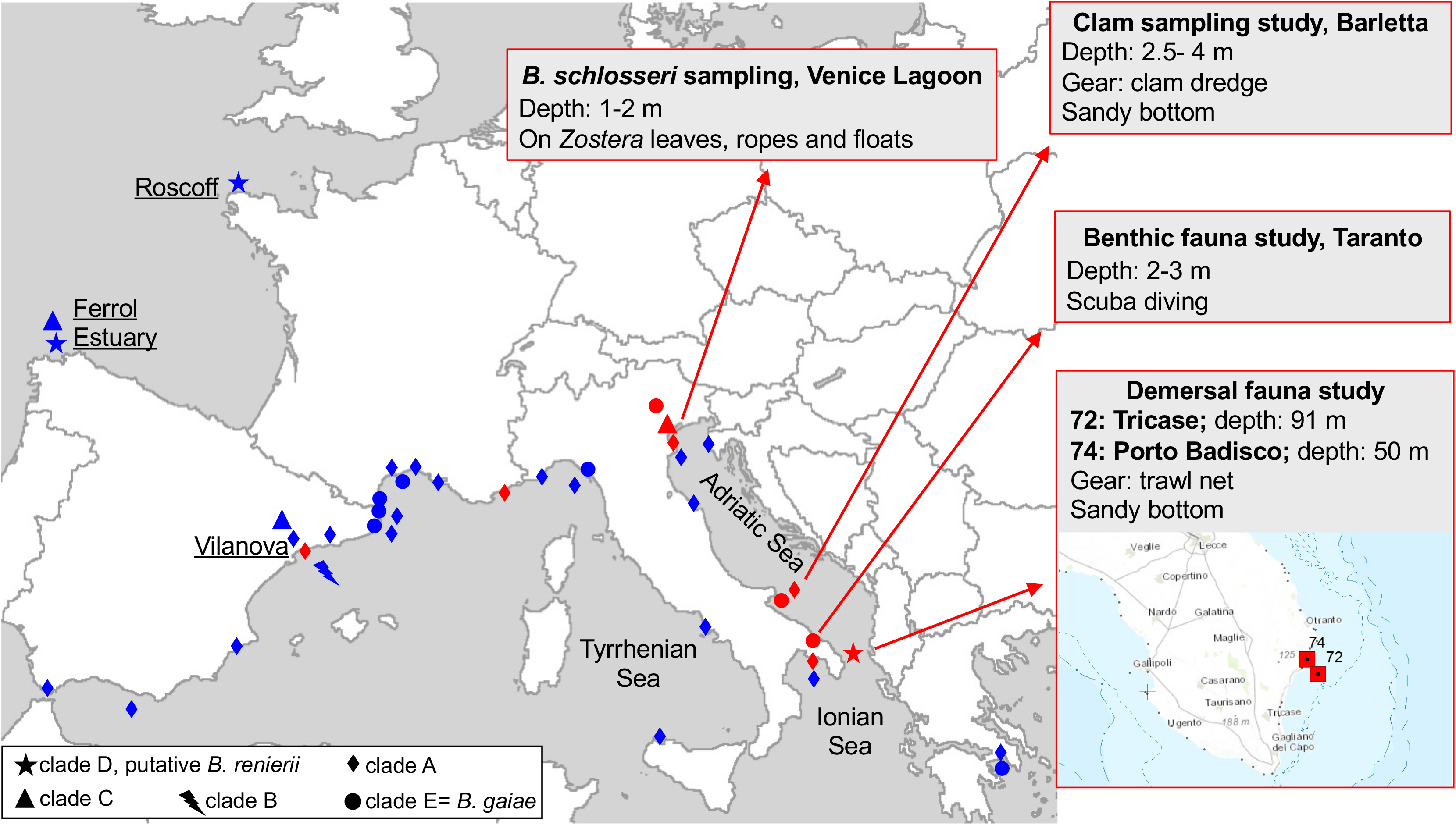
Distribution of the former clades of *B. schlosseri* species complex in the Mediterranean Sea, and presence of the rare clades along the European Atlantic coasts. Red: this study, with localities and dates of our samplings detailed in Supplementary Table S1; blue: data from literature ^9,18,19,32,33^. Notes: in Reem et al.^33^ clade E is reported as clade IV, while clade A includes the clades I, II, and III; localities Fornelos and Ferrol cited in Lopez-Legentil ^18^ and Pérez-Portela ^19^, respectively, are here reported as “Ferrol Estuary”, according to the data provided by the authors (Turon X, personal communication). Indeed, Fornelos is one of the marinas in the city of Ferrol, inside the Ría de Ferrol, while Ferrol is a natural environment at the entrance of the estuary. The map was created with ArcView GIS 3.2 (https://www.esri.com/).

## DISCUSSION

### New primers and amplification strategy

We have here set new primers and a nested amplification strategy for the amplification of a 860-COI fragment in ascidians. The reverse primers have been designed about 200 bp downstream of the HCO2198 barcoding primer ^22^, while the forward primers have been designed in the same region of the LCO1490 barcoding primer (Figure 1). The reason for the design of new primers was two-fold: the need to have better primers and the attempt to obtain a sequence showing higher phylogenetic signal. The effectiveness of the Folmer’s primers in ascidians is not very high, as testified by the bacterial and other contaminant sequences sometimes amplified from ascidian samples (see for example the AY116597 and AJ830012 entries in ^67^ and ^68^). Also based on our experience, most ascidian samples failed to amplify using the Folmer’s primers. The difficulties encountered with the Folmer’s primers have also led researchers to design alternative primers specific for certain ascidian families, genera or species ^23–28^. These alternative primers have even been designed for botryllids and other styelids: see the BvCOIF/BvCOIR primers for *Botrylloides violaceus* ^69^, BsCOIR for some *B. schlosseri* specimens ^9^, and Ah-COIF/Ah-COIR for *Asterocarpa humilis* ^70^. In most cases, the alternative forward primers have been positioned downstream the LCO1490 Folmer’s forward primer, in order to take advantage of the ascidian COI sequences already available, or at exactly the same position of LCO1490 but integrating nucleotide changes or degeneracies (see some examples in Figure 1-B). However, none of them has shown a successful rate of amplification in a wide ascidian range such as to replace the usage of the Folmer’s primers. Even the degenerate primer jglco1490, an upgrade of LCO1490 designed to be able to match the major marine invertebrate groups, has shown low successful rate in ascidians ^71^, probably because it does not consider the possible mismatch at the penultimate 3’ end position (Figure 1-B). As novelty, our dinF primer has been designed with boundaries slightly shifted upstream the position of LCO1490, in order to take into account the peculiarity of the mt genetic code (Figure 1-B). Moreover, the reverse primers have been positioned so as to amplify an additional (200 bp long) region at the 3’ of COI. The extension of the COI fragment also agrees with what observed in diploblasts, where sequencing at the 3’ end of the Folmer’s fragment has been found to be advantageous for species identification and population studies ^72^. Remarkably, our primers and PCR strategy work well not only in botryllids (Table 3) but also in numerous non-botryllid ascidians (Table 2 and Supplementary Table S2), thus strongly suggesting that our approach could be widely used in other ascidian groups. Indeed, only seven out of 55 tested specimens failed to be amplified with our new primers.

### Species delimitation outside clade A

Both ABGD and bPTP species delimitation analyses performed on 860-COI gave the same results, with the only exception of clade A, which is discussed in the next section. These analyses identify as distinct OTUs (*i.e.,* different species) all known species, and all previously identified cryptic species (see yellow diamonds in Figure 2). The OTUs containing more than one sequence are also recognized as highly supported clades in the 860-COI phylogenetic reconstructions (see clades A, clade C, *B. gaiae* and the putative *B. renierii* in Figure 2). More importantly, the species delimitation analyses have led to some unexpected results. This holds true for the five OTUs identified among the nine specimens initially recorded at the Queensland Museum just as *Botrylloides* sp. (Table 3, Figure 2, and Supplementary Table S1). Thanks to the availability of the corresponding formalin-preserved material, these OTUs have been confirmed to be different species. Indeed, their morphological analysis has demonstrated that they correspond to one undescribed species and four old Herdman’s species (*i.e., B. leptum*, *B. jacksonianum, B. pannosum* and *B. anceps)*. The latter two have been referred as “*confer*” (cf.) due to some degree of uncertainty that could only be solved by comparing our specimens to the relative type materials. The re-description of the original species of Herdman ^73,74^ and the description of the new species are beyond the scope of this work and will be published elsewhere. However, these results reveal a botryllid diversity in the Indo-Pacific area higher than expected on the basis of the currently accepted taxonomy. Indeed, the synonymies of the three species *B. leptum*, *B. pannosum* and *B. jacksonianum* with *B. leachii,* proposed by Kott ^75,76^ and followed also by the WoRMS database (http://www.marinespecies.org/aphia.php?p=taxdetails&id=250081), appear to be invalid. Therefore, our results support the resurrection of old species, since they identify as distinct lineages species previously sunk in synonymy and open the possibility for the description of at least one new species.

Another unexpected result concerns *B. primigenus,* whose two analysed specimens are consistently identified as two distinct OTUs, *i.e.,* as two candidate species, by both ABGD and bPTP (Figure 2). Unfortunately, the lack of a formalin-preserved fragment for each of these specimens makes impossible the morphological re-analyses indispensable to solve this issue. Possible explanations of this result could be an erroneous original taxonomic assignment of the specimens or an unexpectedly high genetic diversity indicative of the existence of cryptic species within *B. primigenus*.

Overall, the 860-COI has revealed the presence in our specimen collection of an unexpected diversity and has allowed focusing attention on peculiar specimens/species toward which to direct further morphological analyses.

### Clade A of the B. schlosseri species complex

The 860-COI strongly supports several new sub-clades within clade A that are left unresolved by the Folmer’s fragment both in our study (Table 4) and in many published trees still based on the Folmer’s fragment ^9,20,57^. Remarkably, in previous publications only the amino acid translation of the Folmer’s fragment has provided a phylogeny almost fully resolved both within and between the *B. schlosseri* cryptic species ^18^

In accordance with the phylogenetic reconstruction results, the species delimitation analyses recognize a different number of OTUs within clade A depending on the sequence length (860-COI, Folmer’s fragment) and the method (initial or recursive ABGD results, bPTP). Indeed, the 860-COI fragment recognizes one or two OTUs in clade A based on the method (see vertical bars in Figure 2), while the Folmer’s fragment identifies a single OTU (yellow diamonds in Figure 3). Moreover, some ABGD recursive partitions of both 860-COI and the Folmer’s fragment split clade A in 2-4 OTUs (depending on the distance metric and the prior intra-specific divergence: see Table 5). In particular, in these recursive partitions, 860-COI recognizes only A sub-clades corresponding to nodes strongly supported by the 860-COI phylogenetic tree (compare Table 5 and Table 4), while the Folmer’s fragment identifies even OTUs outside clade A that are clearly unreliable (see the multiple OTUs of *B. leachii* and *B. gaiae* in Table 5). Thus, at least for the ABGD recursive partitions, 860-COI seems to provide more trustworthy results than the Folmer’s fragment. Finally, it is remarkable that when clade A is split in multiple OTUs, these OTUs always include the sub-clade A1, *i.e.* one of the strongly supported clade in the 860-COI phylogenetic tree (see Tables 4-5 and Figure 2).

Taking into account all these results, it is evident that there is currently no conclusive answer to the issue of OTU number/delimitation within clade A. However, it is safe to say that the observed inconsistencies point to a genetic diversity and OTU boundaries deserving further investigations. Of note, this result agrees with two previous studies proposing the existence of cryptic species or ongoing speciation events within clade A, based on the comparison of the entire mtDNA in several intra-species and congeneric ascidian pairs (including the four VE, TR, EA and sc6ab specimens examined also here, and belonging to clade A) ^31,38^. Specifically these two studies recognized three highly divergent groups/specimens within clade A, whose mtDNA nonsynonymous substitution rates are in between those observed in unequivocal intra-species pairs and those found in congeneric ascidian pairs ^31,38^. Remarkably, the multiple OTUs identified within clade A by 860-COI correspond to the three mtDNA-defined groups/specimens. In particular, the sub-clades A1, A3 and A2/A2b include the groups/specimens EA+sc6ab, TR, and VE, respectively, defined by the mtDNA ^31^. Thus, it is important to note that the 860-COI alone allows detecting the same OTUs previously identified based on the entire mtDNA, supporting the view that this marker could be very useful for the identification of OTUs and cryptic species in other ascidian species complex.

Considering the clade A morphological complexity and worldwide distribution, we believe that a conclusive answer to the issue of OTU number/delimitation in this clade can only be provided by applying an integrative taxonomy approach to all identified OTUs/sub-clades of clade A. Future studies should analyse a larger number of specimens and combine information from morphological, ecological and life history traits with the analysis of multiple reliable molecular markers. The resolution of this matter is crucial since *B. schlosseri* is a model organism widely used for investigating several important processes in basal chordates, (such as regeneration, stem cells migration, apoptosis and allorecognition, etc) ^3,5,6^ and it is also a widespread marine invaders ^9,34–36^. It should not be forgotten that the recent morphological re-description of *B. schlosseri* has designed as neotype a COI haplotype corresponding to the VE, NeoB and NeoC specimens ^39^, *i.e.*, a haplotype belonging to the here-defined OTU/sub-clade A2/A2b (see Figure 2 and Table 5). On the opposite, the genome sequencing project of *B. schlosseri* was performed on the sc6ab specimen ^66^, thus on a member of the A1 OTU/sub-clade (Figure 2). Thus, the possible existence of cryptic species within clade A would have substantial effects on the definition and delimitation of this model organism, and needs to be addressed using an exhaustive approach in the future.

### COI as phylogenetic marker in Botryllinae

Although the botryllid phylogeny was not the aim of this research, phylogenetic trees were here reconstructed as an essential requirement for the tree-based coalescence method of species delimitation (*i.e.*, PTP) and as a general method of sequence comparison. Therefore, we can take advantage of these molecular phylogenetic reconstructions to evaluate and compare the resolving power of both the 860-COI and the Folmer’s fragment as phylogenetic markers in the subfamily Botryllinae. Indeed, it should be noted that, so far, the ascidian phylogenies analysing the largest number of species have been carried out using COI and/or the 18S rDNA ^19,77–79^.

The low phylogenetic resolution of the COI Folmer’s fragment in ascidians was already observed at order ^78^ and at family level, including within Styelidae ^19^, but was not investigated at subfamily level. Our data show that the Folmer’s fragment has a low phylogenetic resolution even at subfamily level (within Botryllinae; see Figure 3), thus even at shortest evolutionary distances. Unfortunately, the elongation of the Folmer’s fragment into the 860-COI does not improve the resolution of the Botryllinae tree (compare Figure 2 to Figure 3). Indeed, all our trees are characterized by the lack of resolution at level of the basal nodes, and by polytomies or insufficient support for most nodes describing the relationships between known species (Figure 2 and Figure 3). Therefore, we can conclude that both these COI fragments have insufficient phylogenetic signal for solving the Botryllinae phylogeny.

The low resolving power of COI observed within Botryllinae is probably due to the very fast evolutionary rate typical of all tunicates ^27,49,77,80–82^. This peculiarity makes COI a good phylogenetic marker within ascidians only at very short evolutionary distances, such as those found within species or between cryptic species ^32,83–88^ (this study), but could cause the lack of phylogenetic signal already at the evolutionary distances observed between most species and within a subfamily (*i.e.*, within Botryllinae).

Overall, these data suggest that the Botryllinae phylogeny can be resolved only through an integrative taxonomy approach that includes the investigation of multiple markers, both nuclear and mitochondrial, and/or wide phylogenomic analyses. Previous attempts to elucidate the Botryllinae phylogeny have focused on Japanese species using both morphological and non-morphological characters (*i.e.,* the 18S rDNA, number of stigmatal rows, brooding organ formation and morphology, allorecognition behaviour, vascular system formation, and life history) ^13,29^. These phylogenies have supported the monophyly of *Botrylloides*, and a possible polyphyly of the genus *Botryllus*. Moreover, they have confidently placed *Botrylloides scalaris* sister to all other analysed botryllids ^13,29^. Our COI phylogenetic trees (Figure 2 and Figure 3) do not allow drawing any conclusions about the mono- or poly-phyletic status of the two genera, however they strongly support the existence of a monophyletic clade consisting of most *Botryllus* species except for *B. scalaris* and *B. primigenus*, whose positions remain unresolved or partially resolved (see the black underlined names of *Botryllus* species in Figure 2-3). Of note, in the Bayesian tree of the 860-COI, *B. scalaris* is one of the earliest diverging taxa together with *Botrylloides conchyliatus* (BPP=0.9 in Figure 2).

### Finding rare clades

Our wide botryllid sampling provides new data on the geographic distribution of two rare cryptic species of the *B. schlosseri* species complex, *i.e.* clade C and D ^69^.

Indeed, for the first time we have found the rare clade C in the Northern Adriatic Sea, *i.e.,* in a region other than the Western Mediterranean and the European Atlantic coasts where it was previously found (see red and blue triangles in Figure 4 and Supplementary Table S1).

Likewise, for the first time we have found the rare clade D, here referred as putative *B. renierii*, in the Mediterranean Sea, while it was previously found only along the European Atlantic coasts (see red and blue stars in Figure 4 Supplementary Table S1). Remarkably, not only the few analysable morphological characters (Supplementary File S1) but also the geographic location and the seabed type of our clade D specimens supports their assignment to *B. renierii*. Indeed, *B. renierii* was firstly recorded in the Northern Adriatic Sea, off the Venetian coasts (type locality), and then in several other Mediterranean localities, where it colonises sandy-muddy bottoms at a depth exceeding 15-20 m by attaching to shell fragments and other small portions of hard substrates ^42,53^. Here, we have found clade D in the Southern Adriatic Sea on sandy bottom at a depth of 50-91 m (Figure 4), *i.e.*, in a locality and on a seabed/depth congruent with those typical of *B. renierii.* Therefore, all these data point to the identification of clade D as *B. renierii*. Its original misidentification as *B. schlosseri* can be easily explained when considering that *B. renierii* was for a long time synonymised with *B. schlosseri* ^89^, and was re-described and recognised as a valid species only in 2011 ^42^.

Regarding clade E, widely distributed in the European waters ^9,32^ and recently described as the new species *B. gaiae* ^38^, we have found it in the Adriatic and Ionian Sea, while in the Mediterranean Sea it was previously found only in the Western region and the Greek Aegean Sea ^33^ (see red and blue circles in Figure 4).

As shown in Figure 4, it is striking that some of the *B. gaiae* and clade D findings come from samplings with methods that explore the sandy and muddy bottoms, *i.e.* environments less frequently colonised by ascidians. Shallow-water ascidians are found mainly on hard substrates, however numerous ascidians, belonging to various families, have evolved peculiar adaptations to habitats characterized by soft loose substrata ^90,91^. Our result leads to speculate that *B. gaiae* and clade D have adapted to colonise or are typical of these habitats. The last hypothesis perfectly fits to clade D, *i.e.*, the putative *B. renierii*, since *B. renierii* is one of the few botryllid species characteristic of sandy bottoms ^42^. The rarity of clade D could be then only illusory and consequent to its fortuitous findings in the commonly monitored habitats and in the hotspots of alien species introduction where it was previously found ^18,69^, that is outside of its typical habitat. These observations also suggest that the monitoring of habitats with soft loose substrata and the usage of more diverse sampling methods, even in previously explored geographical areas, could help identify a highest ascidian diversity.

## CONCLUSIONS

We have successfully amplified an elongated COI fragment, 860-COI, in all 177 analysed *Botryllus* and *Botrylloides* colonies. Our specimen collection was very heterogeneous for sampling localities (from Europe to Australia, Japan, California and Brazil) and source (from archival specimens of museum institutions, to colonies kindly provided by ascidian specialists, up to *ad hoc* or fortuitous samplings), and included also colonies with uncertain taxonomic assignment or subjected to only a gross morphological examination. The 860-COI has allowed us not only to effectively discriminate already described (*i.e.,* known) and cryptic species but also to identify undescribed species, to suggest the resurrection of three old species currently synonymized with *B. leachii* (*i.e.*, *B. leptum*, *B. pannosum* and *B. jacksonianum*), and to propose the assignment of clade D to *B. renierii*. Moreover, the 860-COI improved our knowledges of the relationships within clade A of *B. schlosseri,* and highlighted the presence in this clade of a “suspected” high genetic diversity, suggestive of the existence of up to three cryptic species. Together with previously published data on clade A, our result underlines the need to further investigate the taxonomic status of this clade according to an integrative taxonomy approach, that is combining several molecular markers to non-molecular characters (morphology, ecology, development, behaviour, etc.). The case of clade A also shows that 860-COI should be preferred to the Folmer’s fragment for detecting cryptic species and for delimiting very closely allied species in ascidians, *i.e.*, in analyses at very short evolutionary distances. Finally, the 860-COI region was successfully amplified in 48/55 non-botryllid ascidians belonging to several families/orders, suggesting that the relative primers should work well in almost all ascidian taxa, and that this fragment could become a standard DNA barcode for ascidians.

## DATA AVAILABILITY

All sequences generated during this study were submitted to the ENA/GenBank nucleotide database. Their Accession numbers are also reported in Supplementary Tables S1–S4

## ACKNOWLEDGEMENTS

This study is dedicated to the memory of Yasunori Saito. We thank the MEDITS survey programme, the Florida Museum of Natural History (USA), the The Steinhardt Museum of Natural History (National Center for Biodiversity Studies, Israel), the Queensland Museum (Australia) and all peoples listed in Supplementary Table S1 and S3 for kindly providing ascidian samples. C.G. acknowledges the support of MIUR, Italy, (PRIN-2015, grant number: 2015NSFHXF); the Molecular Biodiversity Laboratory of the Italian node of Lifewatch (CNR, Consiglio Nazionale delle Ricerche) and the University of Milano (Italy). D.H. acknowledges the support of the Israel Science Foundation (ISF) grant No. 161/15.

## AUTHOR CONTRIBUTIONS

C.G., D.H. and F.G. designed and planned the study. M.S., M.H-S., M.MN., F.Mo., and C.G. performed PCR experiments. M.E., F.Mo. and F.Ma. performed morphological analyses and wrote the relative part of the manuscript. C.G. and M.S. carried out sequence comparisons and evolutionary analyses. M.S. and C.G. drafted the initial manuscript. All authors revised the manuscript and contributed to data interpretation. C.G., D.H., and F.G. finalized the manuscript. All authors approved the submitted version.

## SUPPLEMENTARY MATERIAL

**Supplementary Table S1**

Botryllinae specimens analysed in this study together with relevant information (specimen and haplotype name, provider, sampling site, collection date, sequenced amplicon, final species assigned, ENA Accession Number).

**Supplementary Table S2**

Complete COI sequences of ascidians analysed in this study, with number of mismatches (mism) to our four new primers and to the two Folmer’s primers.

MP= Mismatch at the Penultimate 3’end position; NW= Not Working due to mispairing at the 3’ terminal position; yellow background: mismatch number > 3. All analysed COI sequences were obtained from entries of complete or almost complete mitochondrial genomes

**Supplementary Table S3**

Origin of the non-botryllid samples and AC number of the 860-COI sequence.

**Supplementary Table S4**

Previously published COI sequences of Botryllinae and *Symplegma* included in the alignment and phylogenetic analyses.

**Supplementary File S1**

Morphological descriptions of the putative *Botryllus renierii* specimens from the Ionian Sea

**Supplementary File S2**

Amplification conditions of the 860-COI fragment using different DNA polymerase

