## Supplementary File S1 for "An elongated COI fragment to discriminate botryllid species and as an improved ascidian DNA barcode"

### *Botryllus* indet.

*Sampling*

Colonies were collected on 6th June 2012 during a demersal fauna study off sandy bottoms of the Ionian Sea (Italian coasts) by using trawling nets. In particular, the collection was performed under the MEDITS survey programme, an International bottom trawl survey, which intends to produce basic information on demersal and benthic species of the Mediterranean Sea in term of population distribution and demographic structure.

Table 1 reports the details of the trawl net haul (a haul comprises a single experimental trawl fishing activity of 30 minutes).

Code station: st.72

Geographic coordination: 40°01'44"N 18°31'04"E

Locality: North-eastern Ionian Sea along the Italian coasts (Tricase), 91 m depth.

Code station: st.74

Geographic coordination: 40°03'51"N 18°29'18"E

Locality: North-eastern Ionian Sea along the Italian coasts (Porto Badisco), 50 m depth.

*Material examined*

Orange colony exam 1; 6/6/12; Medits; st.72

Orange colony exam 2; 6/6/12; Medits; st.72

Violet colony exam 1; 6/6/12; Medits; st.74

Violet colony exam 2; 6/6/12; Medits; st.74

*Description*

The material examined was mostly degraded except for very few zooids/portions of the colonies that appear in better conditions.
The colonies were cushion and globular in shape and characterized by a very thick matrix (about 3-4 cm), without any sand embedded (Fig. 1 e). The zooids were arranged in star-shaped or oval-shaped systems with a common cloaca (Fig 1 a). They were positioned vertically with respect to the colony surface (Fig. 1 d), almost cylindrical in shape and about 5 mm in length (Fig 1 c). Several spherical ampullae were evident within the vascular network of the matrix as well as among the systems on the colony surface (Fig. 1 b). The branchial sac of the zooids showed more than 10 rows of stigmata. The stomach was placed partially below to the branchial sac (Fig. 1 c). On the mesial side are visible 8 spiral folds (Fig. 1 g-h), while the parietal side of the stomach was not clearly distinguished (Fig. 1 g-h). Above the gut loop were clearly visible the mature testes, fan-shaped and deeply divided (Fig 1 c-i). The above-described features reveal that these colonies are not *B. schlosseri sensu* the recent re-description of the species by Brunetti *et al.* (2017). Indeed *B. schlosseri* shows a minor number of rows of stigmata (7-8) and a clearly encrusting colony.

*Results and discussion*

The bad condition of the material suggests the application of a precautionary approach and a consequent more conservative identification as *Botryllus* indet. Indeed, according to the Open Nomenclature qualifiers (Sigovini *et al.,* 2016), "indet." means that the specimen is indeterminable beyond a certain taxonomic level due to the deterioration or lack of diagnostic characters. However, some characteristics of these specimens suggest that they can be assigned to the species belonging to the old genus *Polycyclus* (nowadays in synonymy with *Botryllus*) and in particular to *B. renierii* as described by Brunetti (2011). These features are: the globular shape of the colonies with a thick matrix, the star or oval arrangement of the systems, the size of the zooids and, as for the internal zooid anatomy, the number of stigmata rows, the shape of the stomach and the shape of the testis.

**Table 1.** Sampling scheme with indications of the geographical subarea, name of the station, date, geographical coordinates of the start and the end of each trawl net haul with relative depth and duration.

| Geographical sub-area | Station | Day | Month | Year | Start time | End | Longitude start | Iniatial depth | End time | Latitutide end | Longitude end | Depth end | Duration (minutes) |
|---|---|---|---|---|---|---|---|---|---|---|---|---|---|
| Western Ionian Sea (19) | 72 | 6 | 6 | 2012 | 08:47 | 40°02’29’’ | 18°31’39’’ | 88 | 09:17 | 40°01’31" | 18°31’33’’ | 92 | 30 |
| Western Ionian Sea (19) | 74 | 6 | 6 | 2012 | 07:27 | 40°04’36’’ | 18°29’25’’ | 59 | 07:57 | 40°03’03’ | 18°29’06" | 39 | 30 |

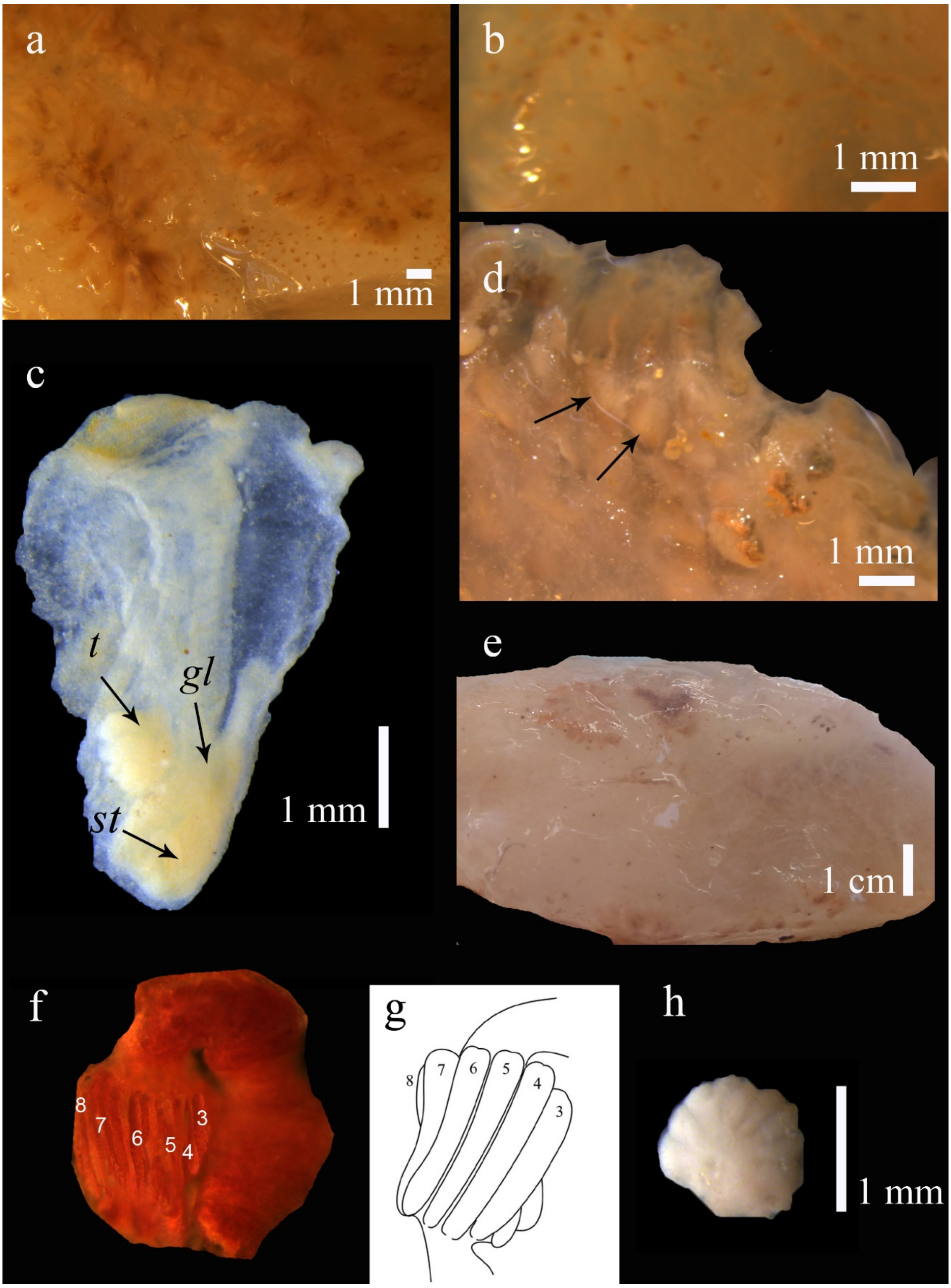


**Figure 1**. *Botryllus* indet. (a) Zooids arranged in oval-shaped systems with several spherical ampullae visible among them. (b) Spherical ampullae present within the matrix vascular network. (c) Zooid with atrial languet clearly visible, testis positioned above the gut loop, stomach lateral. *t,* testis; *gl,* gut loop; *st,* stomach. (d) Zooids positioned perpendicular to the surface of the colony; (e) Cushion shaped colony with very thick matrix. (f-g) Comparison between the mesial side of the stomach of the collected colonies (on the left, f) with the same feature described for *B. renierii* by Brunetti (2011) (on the right, g). (h) Fan-shaped testis, deeply divided.
