## Supplementary File S2 for "An elongated COI fragment to discriminate botryllid species and as an improved ascidian DNA barcode"

Amplifications of Botryllinae with the high fidelity PrimeStar HS or PrimeStar GXL DNA polymerases (TaKaRa Bio Inc.) were performed in a final reaction volume of 25 µl containing: 1X reaction buffer with 1 mM final concentration of $MgCl_2$ (TaKaRa Bio Inc.), 0.2 mM of each dNTP, 0.3 µM of each primer, 1.25 Units of PrimeStar HS or GXL DNA polymerase (TaKaRa Bio Inc.) and 20-200 ng DNA. Amplification conditions were: 30 cycles with denaturation for 10 s at 98°C, annealing for 15 s at 44-50°C (depending on the sample), extension for 1 min 30 s at 72°C; a final elongation step of 5 min at 72°C for the PrimeStar HS and at 68°C for the PrimeStar GXL.

PCRs with the DreamTaq polymerase (Thermo Fisher Scientific) were performed in a final volume of 25 µl containing: 1X reaction buffer with 2 mM final concentration of $MgCl_2$ (Thermo Fisher Scientific), 0.2 mM of each dNTP, 0.4 µM of each of the two primers, 1.25 Units of DreamTaq polymerase (Thermo Fisher Scientific) and 20-200 ng DNA. The amplification conditions were: an initial denaturation for 3 min at 95°C, then 30 amplification cycles (denaturation for 30 s at 95°C; annealing for 30 s at 48-52°C; extension for 1 min 30 s at 72 °C) followed by a final elongation step of 5 min at 72°C.

The obtained amplicons were purified with the Amicon Ultra-0.5 mL centrifugal filter devices (NMWL of 30kDa, Millipore) or the DNA Clean&Concentrator kit (Zymo Research) and directly sequenced according to the Sanger method at the Eurofins Genomics (Ebersberg, Germany) or Microsynth AG (Switzerland). Sequence quality check, assembly, and comparisons were carried out with Geneious ver. 5.5.7.2 (Kearse et al., 2012) .

The dinF/Nux1R fragment of the QM_G335164 specimen was amplified with the EmeraldAmp Max HS PCR Master Mix (TaKaRa Bio Inc.), using 12.5 µl of PCR Master Mix 2X and a final concentration of 0.4 µM for each primer. The amplification conditions were: 30 cycles with denaturation for 10 s at 98°C, annealing for 15 s at 48°C and extension for 1 min 30 s at 72 °C; a final elongation step of 5 min at 72°C. The unpurified PCR product was sent to Macrogen Inc. (Seoul, South Korea), cleaned, and then sequenced on an ABI3730XL sequencer via the Standard-Seq service.

PCRs of 15 non-botryllid were conducted with the Ex Taq (TaKaRa Bio Inc) DNA polymerase. The reactions were performed in a volume of 25 µl of the following composition: 1X reaction buffer with 2 mM final concentration of $MgCl_2$, 0.5M of betaine, 2.5% DMSO, 0.2 mM of each dNTP, 0.5 µM of each primer, 1 Units of Ex Taq (TaKaRa Bio Inc) and 20-200 ng DNA. The amplification conditions were: an initial denaturation for 4 min at 95°C, then 35 amplification cycles (denaturation for 45 s at 94°C; annealing for 45 s at 50°C; extension for 1 min at 72 °C) followed by a final elongation step of 5 min at 72°C. The purification of the amplicons was done using ExoSAP-IT (Thermo Fisher Scientific) following the manufacturer protocol. Sequencing was performed on both strands on an ABI 3500xl Genetic Analyzer (Applied Biosystems) by The DNA Sequencing Unit at the G.S. Wise Faculty of Life Sciences (Tel-Aviv University).
